# Balancing true and false detection of intermittent sensory targets by adjusting the inputs to the evidence accumulation process

**DOI:** 10.1101/2022.09.01.505650

**Authors:** Anna C. Geuzebroek, Hannah Craddock, Redmond G. O’Connell, Simon P. Kelly

## Abstract

Decisions about noisy stimuli are widely understood to be made by accumulating evidence up to a decision bound that can be adjusted according to task demands. However, relatively little is known about how such mechanisms operate in continuous monitoring contexts requiring intermittent target detection. Here, we examined neural decision processes underlying detection of 1-second coherence-targets within continuous random dot motion, and how they are adjusted across contexts with Weak, Strong, or randomly Mixed weak/strong targets. Our prediction was that decision bounds would be set lower when weak targets are more prevalent. Behavioural hit and false alarm rate patterns were consistent with this, and were well-captured by a bound-adjustable leaky accumulator model. However, beta-band EEG signatures of motor preparation contradicted this, instead indicating lower bounds in the Strong-target context. We thus tested two alternative models in which decision bound dynamics were constrained directly by Beta measurements, respectively featuring leaky accumulation with adjustable leak, and non-leaky accumulation of evidence referenced to an adjustable sensory-level criterion. We found that the latter model best explained both behaviour and neural dynamics, highlighting novel means of decision policy regulation and the importance of neurally-informed modelling.

## Introduction

The way that we form decisions is highly adaptable to task demands. It is now widely understood that making accurate decisions on noisy sensory information involves the accumulation of evidence up to a criterion amount or “bound” (***Gold and Shadlen, 2007***; ***Ratcliff and Smith, 2004***). This bound is seen as the primary means to strategically adapt perceptual decision policies to meet various task demands. Significant behavioural and neural evidence has now amassed to establish that bound adjustments are indeed carried out for certain, well-studied scenarios, such as when trading speed for accuracy in discrete, 2-alternative forced choice decisions (***Bogacz et al., 2010***; ***Palmer et al., 2005***), but it is not clear whether this mechanism generalises to all situations that require policy adaptations. In particular, in the pervasive daily life scenario in which unpredictably-timed targets must be detected within continuous streams of noisy sensory information, one’s decision policy must be tailored to rapidly detect as many targets as possible while keeping false alarms to a minimum (***Lasley and Cohn, 1981***). Here, an overly liberal policy could result in an excessive amount of false alarms driven solely by noise in between targets, while an overly conservative policy will miss too many targets (***Gold and Stocker, 2017***; ***Ossmy et al., 2013***). In such contexts, expected target strength is a key factor in setting a decision policy to optimise performance -contexts with weaker targets call for a more liberal decision policy. Although bound adjustments present a natural mechanism for these policy adjustments, this has never been experimentally tested for this scenario.

Much has been learned about the neural implementation of decision bound adjustments from studies of the neural basis of the speed-accuracy tradeoff. In these studies, motor-selective cortical networks have been shown to implement a static increase of evidence-independent baseline activity, emphasising speed by effectively decreasing the amount of cumulative evidence needed to reach the decision-bound (***Forstmann et al., 2008, 2010***; ***Ivanoff et al., 2008***; ***Van Veen et al., 2008***; ***Wenzlaff et al., 2011***). Additionally, recent behavioural and neurophysiological findings have established the operation of a time-dependent “urgency” signal in these motor circuits, an evidence-independent buildup component that drives the decision process closer to a fixed decision-threshold, gradually decreasing the cumulative evidence required to respond (***Churchland et al., 2008***; ***Hanks et al., 2014***; ***Murphy et al., 2016***; ***Thura and Cisek, 2016***). However, little is known about how such mechanisms may participate in continuous detection scenarios, or in the adjustments needed to account for other task-related factors such as the expected target evidence strength. Our initial prediction for the present study was that the decision bound -the criterion amount of cumulative evidence-would be lower in continuous detection contexts containing weaker targets. However, in continuous detection where target onsets are unpredictable, the decision process must operate continuously, offering other parameters that could alternatively be adjusted. For example, if accumulation proceeds continuously it is unlikely to be perfect because this generates excessive false alarms (***Ossmy et al., 2013***) and would instead entail some form of information loss such as through non-linear or leaky accumulation, which has been shown to form a critical adjustable component governing decision-making policies in volatile environments (for review see ***Gold and Stocker, 2017***; ***Glaze et al., 2015***; ***Murphy et al., 2021***).

Our findings indicate that observers can adjust their decision-making policy based on their knowledge of the difficulty context, but not by adjusting bound settings as found for speed/accuracy adjustments. While the electrophysiological data did show both static, global context-dependent adjustments and a growing evidence-independent urgency signal through the inter-target interval (ITI), this urgency component was not set higher in order to reduce the bound in Weak target contexts as predicted. Instead, the bound was reduced in the Strong context, as indicated by a smaller excursion between baseline motor activity and the level reached at the time of movement execution reflecting the action-triggering threshold. We therefore hypothesised that an alternative way to set a conservative decision policy is to reduce the information gathered in the accumulator, and tested two ways of implementing this: 1) leaky accumulation with context-dependent leak and 2) a non-leaky accumulation of evidence referenced to a context-dependent sensory criterion. Fitting these models with bound dynamics directly constrained by the observed motor preparation signals, we found that the latter model produced a superior fit to both behavioural data and the dynamics of a motor-independent signature of evidence accumulation, the centro-parietal positivity (CPP; ***Kelly and O’Connell, 2013***; ***O’Connell et al., 2012***). We are thus able to falsify the initially predicted bound adjustment model in favour of an account based on the adjustment of a sensory-level criterion that regulates the transfer of incoming evidence into the accumulation process.

## Results

### Behavioural signatures of contextual adjustments

We analysed the data of fourteen participants that continuously monitored a cloud of randomly moving dots for unpredictable intermittent targets defined by coherent upward motion. Within each block, the Difficulty Context was set to be Fixed with 1) only Weak targets, 2) only Strong targets, or 3) Mixed with both randomly occurring weak and strong targets, with the latter included to control for differences due to target motion coherence itself as distinct from context (see Fig. 1A). As expected, participants detected targets more accurately and faster when the targets were Strong for both the Fixed and Mixed context (Fig 1 B; main effect of Motion coherence on hit rates in *GMM, χ*^2^(1) = 8.5, p <0.001, and RT in *rANOVA*, F(1,13) = 371.8, p <0.001). Additionally, participants were faster in the Mixed context than the Fixed ones (main effect of Difficulty Context on RT in *rANOVA*, F(1, 13) = 21.3, p <0.001).

**Figure 1.**
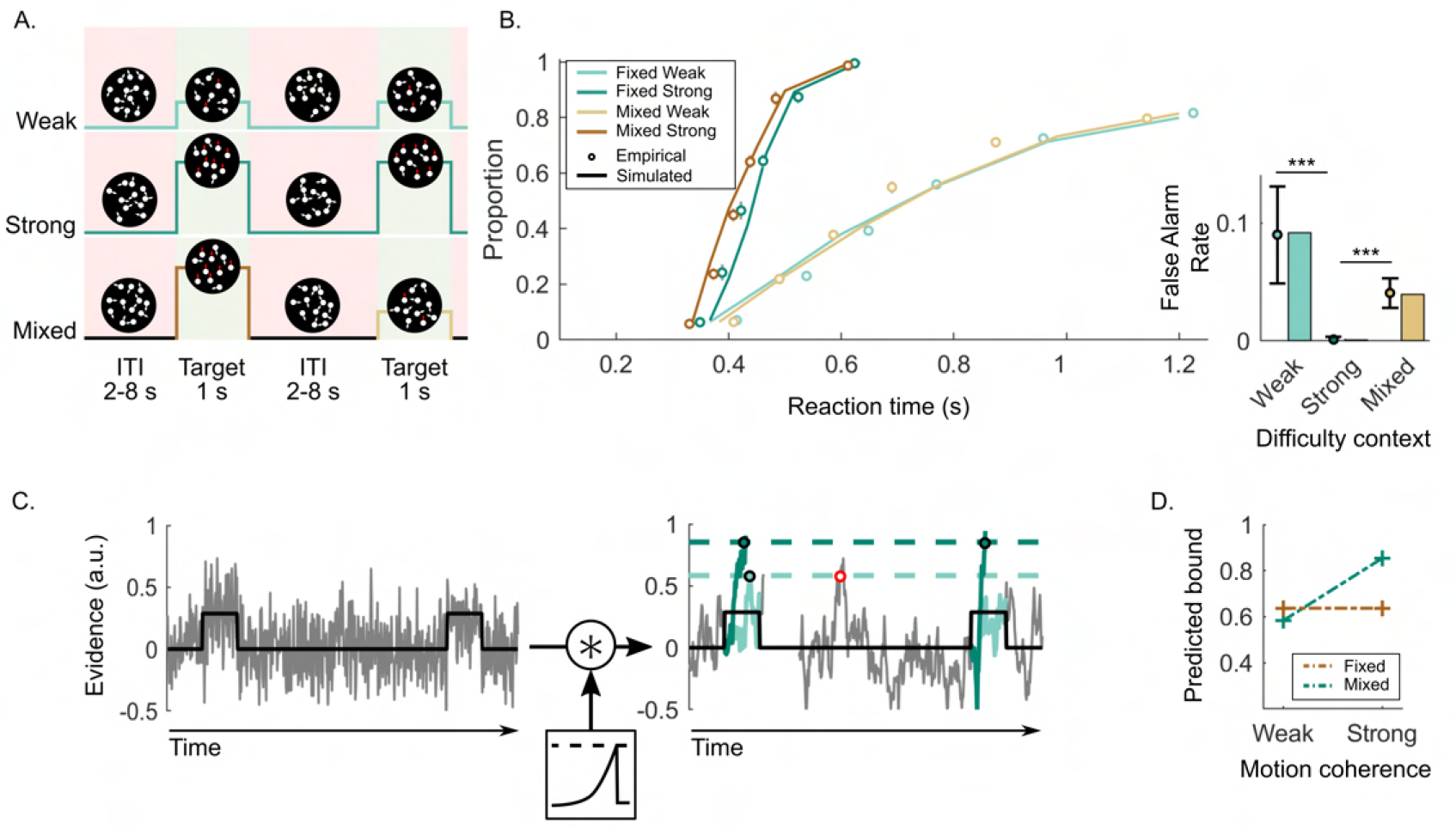
Continuous Random Dot Motion (RDM) detection task and behaviour. **(A)** Participants continuously monitored a cloud of moving dots to detect targets defined by a step change to coherent upward motion for 1 sec. The inter-target interval (ITI) duration varied between 2, 4, 6 or 8 sec. In each block of 24 targets, participants would perform a Weak Fixed Context condition (step increase of 25%), a Strong Fixed Context condition (step increase of 70%), or a Mixed Context condition (25% and 70% equally likely). **(B)** RT cumulative quantile probability functions and the proportions of 2-sec ITI periods containing a False Alarm as a function of Difficulty Context (Dots). Each plot includes simulated data from a fitted leaky accumulator model with bound-adjustment (solid lines in cumulative quantile probability plots and bars). **(C)** Schematic representation of the leaky accumulator model with adjustable bound parameters. Noisy evidence, which steps up during targets, is accumulated until it reaches a context dependent bound. The degree of leakage is a free parameter but constrained to be equal (not adjusted) across contexts. When the target evidence is weak, a more liberal (lower) bound needs to be set to avoid misses; this comes however at the cost of more false alarms (example indicated in red). **(D)** Bound parameter values estimated by the best fitting bound-adjustment model. Error bars represent the 95% confidence interval. P-values are indicated by the asterisks;*p < 0.05, **p < 0.01, and ***p < 0.001.

False alarm rates demonstrate a qualitative signature of contextual adaptations, with approximately zero in the Strong context, higher false alarms rates in the Weak context (0.09 per 2-sec ITI period) and intermediate rates in the Mixed context (0.04 per 2-sec ITI period; Fig. 1B; *GMM, χ*^2^(1) = 5.7, p < 0.001 and *χ*^2^(1) = 5.5, p < 0.001, respectively). In accumulator models of decision making, these patterns can be captured by setting a higher bound when the task is easy, to avoid false alarms while still detecting all targets (see Fig. 1C). Confirming this, a leaky accumulator model with a higher bound for the Strong context (Fig. 1D and Table 1 for all the fitted parameters), is able to accurately reproduce behavioural data, particularly the important context-dependent false alarm pattern (see bars of the false alarm plot in Fig. 1B;*G*^2^ = 16).

**Table 1.**
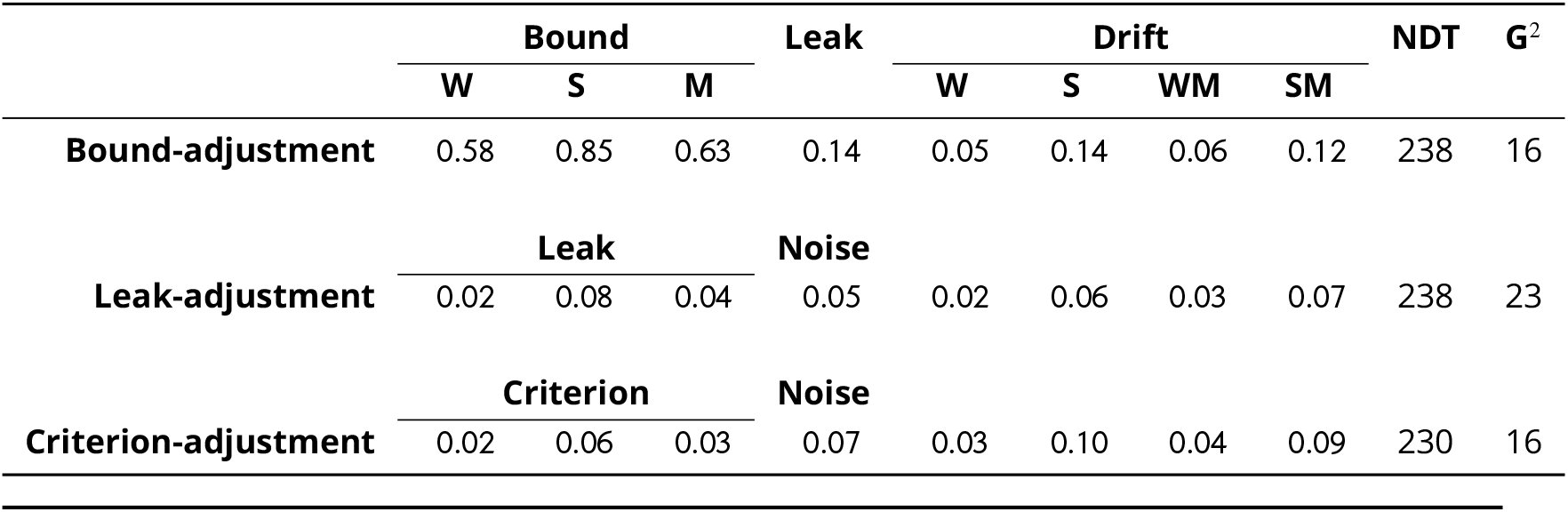
Parameter values estimated for the leaky accumulator model with adjustable bound fit to behaviour alone, and the neurally-informed leaky accumulator model with adjustable leak and non-leaky integration model with adjustable sensory criterion. Goodness-of-fit is provided in the form of G2 metric; note all models have an equal number of free parameters. Abbreviations: NDT is non-decision time, W is Weak Condition, S is Strong Condition, M is Mixed Condition.

### EEG signatures of decision formation: adjustments to Difficulty Context

To test for electrophysiological evidence for the putative bound adjustment, we examined two signals known to reflect the dynamics of decision formation: decreases in Beta frequency band (15-30 Hz) over motor cortex contralateral to the movement, reflecting motor preparation (see Fig. 2A), and the CPP, reflecting motor-independent evidence accumulation (see Fig. 2D).

**Figure 2.**
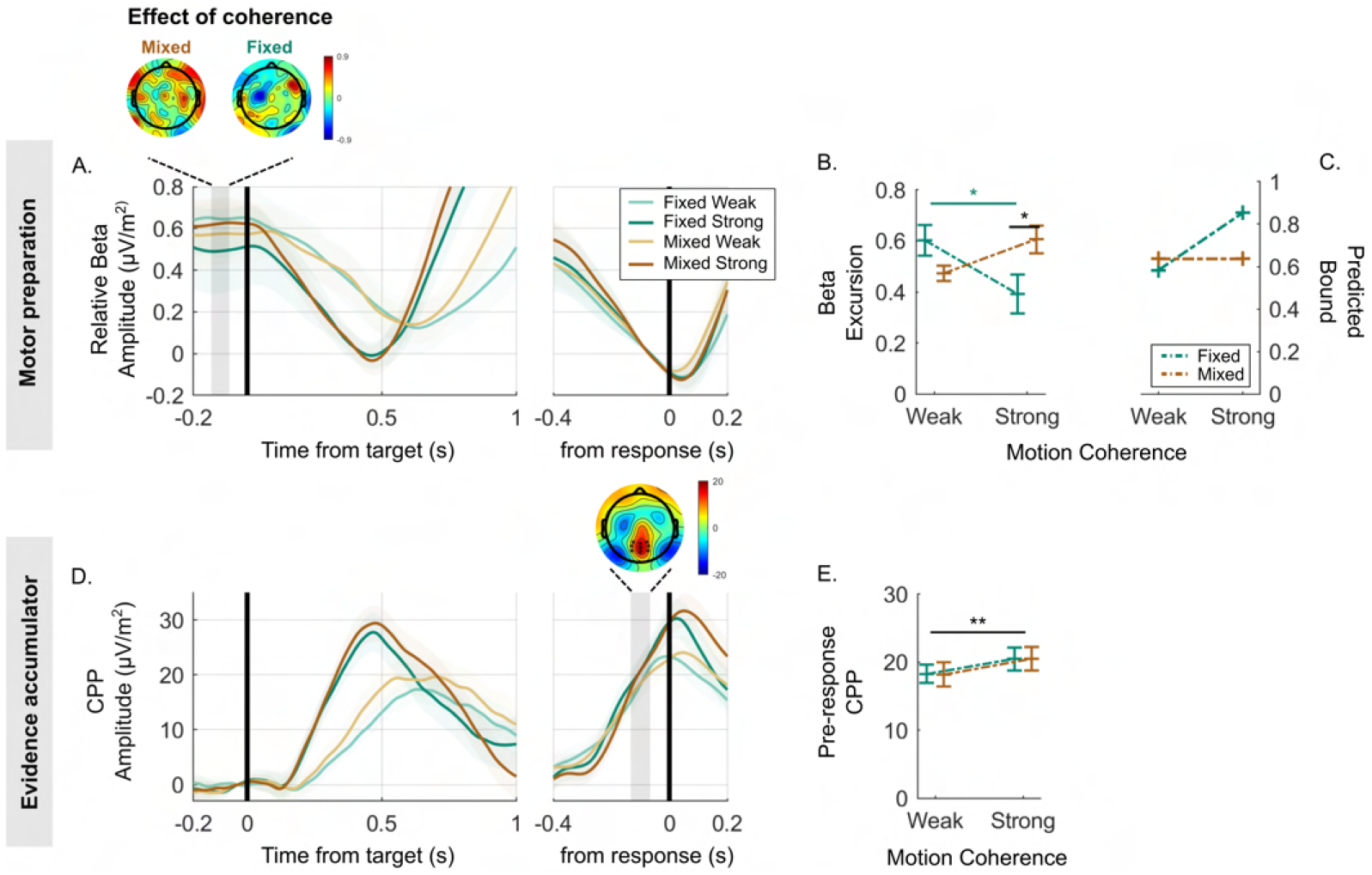
Decreases in Beta amplitude track motor preparation and reveals a lower decision bound for the Strong context. **(A)** Beta amplitude (15-30 Hz) over the motor cortex, aligned to the target (left) and to the response (right), averaged for each condition and expressed as a difference relative to the corresponding pre-response level taken to reflect motor execution threshold (i.e. threshold level is indicated by zero). The scalp topography shows the distribution of the difference between weak and strong targets in the baseline amplitude (again relative to the motor execution threshold). This highlights that the difference in Beta excursion in the Fixed Strong relative to Fixed Weak context is maximal over the motor cortex. **(B)** The average difference of baseline beta amplitude minus pre-response beta amplitude for all conditions, reflecting the ‘excursion’ from starting level to threshold and hence the neural index of bound settings. **(C)** The bound parameter values estimated in the behavioural data fit of the leaky accumulator model with adjustable bound, reproduced from Fig. 1D, showing an obvious qualitative difference. **(D)** CPP signals averaged for each condition, target-locked (left) and response-locked (right). The topography of the indicated pre-response time window shows the cluster of electrodes used for plotting average waveforms. **(E)** Average pre-response CPP amplitude for each condition. Error bars for all plots represent the 95% confidence interval. P-values are indicated by the asterisks;*p < 0.05, **p < 0.01, and ***p < 0.001. **Figure 2—figure supplement 1.** Additional analysis of posterior N2 component as distinct, lower-level process functioning as a relevant transition detector.

Previous studies have shown that Beta oscillatory activity over motor cortical EEG sites is gradually suppressed as participants prepare to make a movement with a contralateral limb (***Donoghue et al., 1998***; ***Pfurtscheller et al., 1996***). This beta-indexed motor preparation builds at an evidence-dependent rate during decision formation and reaches a stereotyped threshold level at the time of motor execution (***Kelly et al., 2021***; ***O’Connell et al., 2012***). Fluctuations in prestimulus Beta activity have been shown to predict within-subject trial-to-trial behavioural variability (***de Lange et al., 2013***; ***Donner et al., 2009***; ***Gould et al., 2012***), and it is systematically shifted closer to its pre-response level under conditions of speed emphasis (***Murphy et al., 2016***; ***Steinemann et al., 2018***; ***Kelly et al., 2021***), underscoring its role as a reliable readout of a decision variable encoded at the motor level.

In line with this body of previous research, we took the amplitude reached by Beta 80 ms prior to response (allowing for motor execution delays) to correspond to the threshold level required to trigger a response in a given condition, and the level in the pre-target baseline period as the decision variable starting point at the time of evidence onset. In this way, the difference between the two, known as the “excursion”, indexes the change in motor preparation required to commit to a detection decision and thus indexes bound settings in a given context (***Heitz and Schall, 2012***; ***Kelly et al., 2021***, see below for separate analysis of baseline and pre-response amplitudes). Difficulty context (Fixed vs. Mixed) and Motion coherence (Weak vs. Strong) had a significant interactive effect on Beta excursion (Fig. 2B; *GMM, χ*^2^(1) = 2.5, p = 0.01). Contrary to the prediction of the bound-adjustment model (Fig. 2C), this was driven by a pattern whereby Beta excursion was smaller, not larger, in the Strong Context relative to other conditions, indicative of a lower decision bound. Within each context, we found that Beta excursion was positively related to RT in all conditions (*z*^2^(1) = −0.12, p = 0.01), indicating that response times are faster on trials where Beta happens to have a pre-target level that is closer to its threshold, further supporting Beta excursion as an index of the decision criterion on cumulative evidence.

More recently, another signature of decision formation was characterised - the CPP - which, like Beta, builds at an evidence dependent rate to a peak around the time of decision commitment and response (***O’Connell et al., 2012***; ***Kelly and O’Connell, 2013***). This signal, unlike Beta, builds even in the absence of motor requirements and does not undergo baseline shifts according to speed/accuracy emphasis (***Steinemann et al., 2018***). The current working hypothesis is thus that the CPP reflects pure cumulative evidence which feeds into motor preparation alongside other evidenceindependent signal components such as urgency. In support of this scheme, simulations of pure cumulative evidence buildup have been shown to match empirical CPP waveforms in their dynamics and pre-response amplitude patterns across conditions and RTs (***Afacan-Seref et al., 2018***; ***Kelly et al., 2021***; ***Twomey et al., 2015***). Here, a hypothetical higher bound setting in the Strong context would predict that more cumulative evidence is required before reaching the movement execution threshold, and hence a higher CPP amplitude. More specifically, a true contextual adjustment effect should give rise to a CPP amplitude difference in the Strong relative to Weak context over and above any difference between high and low coherence targets in the Mixed context. This however is not observed in the empirical pre-response CPP, where only a significant Motion Coherence effect was found (*GMM*: *χ*^2^(1) = 5, p < 0.001; Figures 2D and E). Tracing the waveforms beyond the pre-response measurement window shows that the CPP tends to climb higher for the Mixed Strong than Fixed Strong condition, if anything (Fig. 2D). The electrophysiological data are therefore inconsistent with the bound-adjustment model, raising the question of whether an alternative mechanism can explain the behavioural markers of policy adjustment while also producing the observed dynamics in neural decision signals.

### Alternative neurally-informed models

Following a recent approach (***Kelly et al., 2021***), we constructed models whose decision bound settings were constrained to match the observed Beta excursion measurements. We use Beta as it best corresponds to the motor-level decision variable which is ultimately subjected to a threshold. After fitting to behaviour, we then examined whether key dynamical features of evidence accumulation predicted by the models matched those of the observed CPP time courses in a further validation step. To determine the Beta activity-based constraints, it was important to first characterise the dynamics of motor preparation during the ITI more fully. Beta waveforms during the full ITIs showed that in addition to the static shift toward the motor execution threshold in the Strong context, there was a strong tendency for Beta to increase over the course of the ITI in all contexts (Fig. 3A lower), which mirrored a tendency for false alarm rate to increase during the ITI (Fig. 3A, *upper*). This is consistent with previous observations of an evidence-independent buildup component in signals related to motor preparation attributed to gradually increasing urgency (***Cisek et al., 2009***; ***Hanks et al., 2014***; ***Murphy et al., 2016***; ***Standage et al., 2011***; ***Thura et al., 2012***; ***Thura and Cisek, 2014, 2016***), seen sometimes in anticipation of evidence (***Corbett et al., 2021***; ***Kelly et al., 2021***). This growing urgency component, which in evidence-accumulation models combines with cumulative evidence in a decision variable building towards a constant action-triggering threshold, is mathematically equivalent to, and a well established neural implementation of, a collapsing decision bound (***Churchland et al., 2008***; ***O’Connell et al., 2018***). Here, the decrease in Beta amplitude relative to its ending ‘threshold’ level reflects a motor preparation signal that grows as a concave nonlinear function from ITI onset (Fig. 3A, *lower*), reducing the distance remaining to the threshold level with time as target evidence is awaited. Both static effects and nonlinear time-dependent urgency are thus reflected in the Beta amplitude in the ITI. To use this to constrain our alternative models, the grand-average Beta amplitude waveform in the 8 sec ITI was taken and normalised according to the motor execution threshold indexed by average pre-response beta in a given condition, and a 2nd-order polynomial was then fitted to capture the smooth urgency trend relative to threshold (see methods for more details). This signal was added to the output of evidence accumulation and the resultant decision variable was subjected to the ultimate, constant decision threshold (see Figures 3B and C).

**Figure 3.**
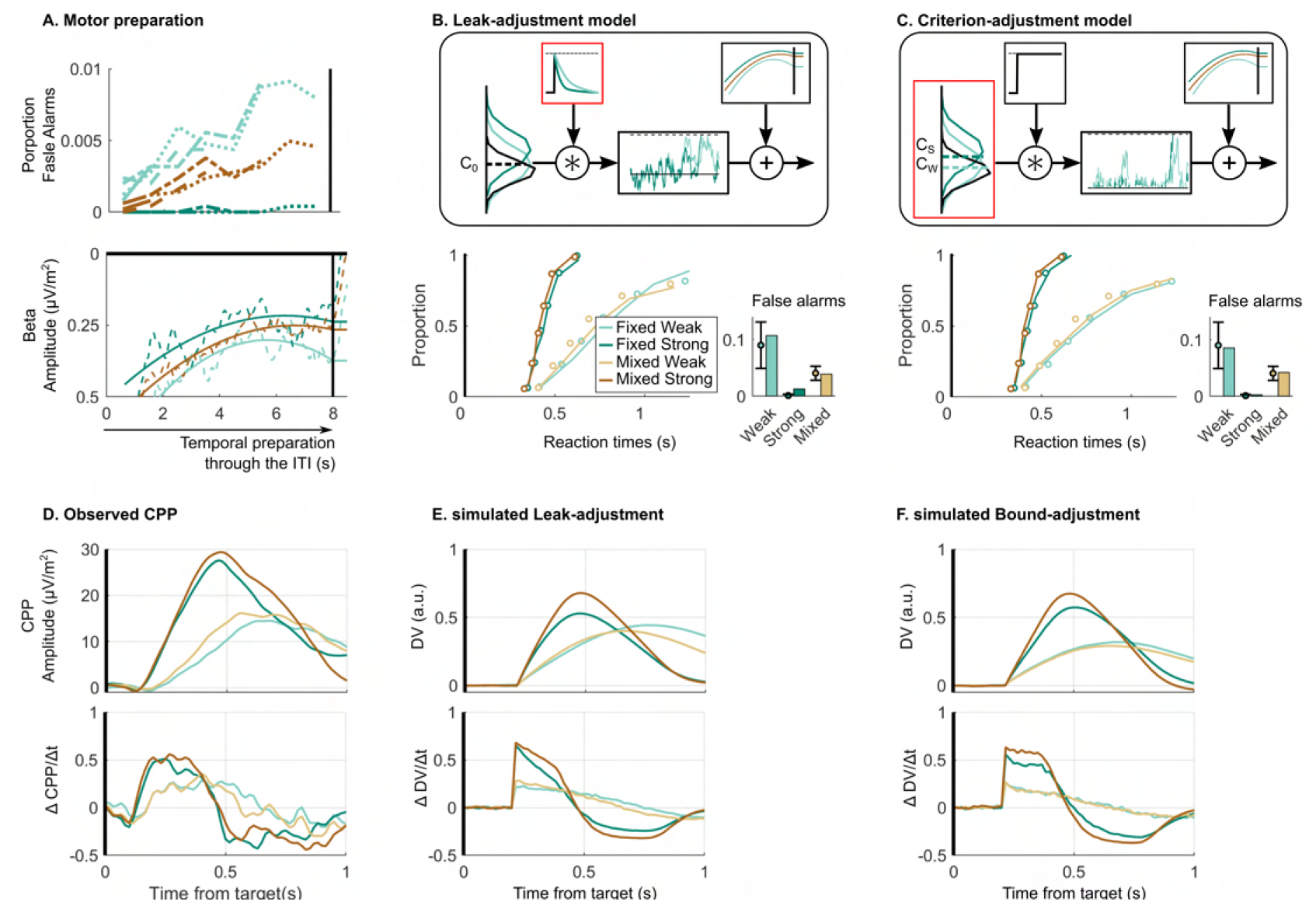
Neurally informed modelling. **(A)** Motor preparation, also termed urgency, throughout the ITI. The proportion of false alarms increases throughout the ITI (*upper*). This corresponds to an increase in motor preparation throughout the ITI reflected in the Beta amplitude (*lower*). Beta amplitude is plotted for the 8 sec ITI normalised relative to the corresponding pre-response level taken to reflect motor execution threshold. The normalised Beta amplitude is used to model temporal expectation through the ITI, which generates the increase of false alarms. Model evaluation for the neurally-informed **(B)** leak adjustment and **(C)** sensory criterion-adjustment model comparing empirical and predicted RT cumulative quantile probability distributions and False Alarm rates (per 2-sec ITI period). Above are model schematics illustrating the different mechanisms of continuous accumulation and context-dependent adjustment. In the leak-adjustment model, evidence is referenced to the centre of the noise distribution (the ‘zero’ point) and the degree of leakage, represented in the time-constant of a convolution kernel, is free to change across contexts. Meanwhile in the criterion-adjustment model, evidence is referenced to a criterion ‘zero’ point that is adjustable across contexts, and fed to a non-leaky accumulator with a lower reflecting bound at zero to preclude negative accumulation. **(D)** Empirical average target-locked CPP signals (upper row) for each condition as well as their first derivative (lower row). This can be compared to the simulated accumulator process for both **(E)** the leak-adjustment and **(F)** criterion-adjustment models. The buildup rate of the CPP does not immediately begin to fall steeply in the way predicted by the leaky accumulator model. In these simulations, CPP is simulated as the cumulative evidence without direct urgency influence (***Kelly et al., 2021***). Without loss of generality for the behavioural responses, we assumed that after reaching commitment, there is a delay of 80 ms before the CPP stops accumulating and falls linearly over 416 ms back to zero, implemented identically in all contexts and in both models. This was based on observed post-response dynamics in the real CPP. **Figure 3—figure supplement 1.** Comparison of response-locked dynamics in the empirical CPP with the simulated pre-response cumulative evidence.

Using these constraints, we explored other possible mechanisms of continuous accumulation and context-dependent adjustment. First, we considered that the same leaky accumulation mechanism as the bound adjustment model may be at play but with the adjustment across contexts being applied to leak rather than bound (Fig. 3B). Here, a more forgetful (higher leak) integration would reduce the risk of false alarms driven by noise (***Glaze et al., 2015***; ***Murphy et al., 2021***; ***Ossmy et al., 2013***) in a way similar to an increased bound, and thus presents an alternative means of setting a conservative policy. Second, we considered that alternatively, a context-dependent evidence criterion could be set at each momentary evidence sample serving as a ‘zero’ reference placed between noise and signal evidence distributions, similar to the “drift criterion” principle proposed by Ratcliff and colleagues (***Ratcliff, 1985***; ***Ratcliff et al., 1999***; ***Ratcliff and Tuerlinckx, 2002***). This criterion is distinct in meaning to that of signal detection theory in that it does not on its own define the ultimate decision rule to categorise signal versus noise, but rather serves as a reference subtracted from the evidence so that only evidence samples substantially above the noise would be positively accumulated towards the decision-bound, while noise alone would conversely tend to count negatively. To prevent runaway negative accumulation in this model, we adopt a reflecting lower bound at zero in the accumulator process (***Usher and McClelland, 2001***). Thus, this last model has two major distinctions: it assumes that an adjustable criterion is set directly on the momentary evidence in addition to the adjustable decision-bound on the cumulative evidence (with the latter measured directly from beta excursion), and involves a different form of information loss than leak which nevertheless has a similar effect of reducing risk of false alarms due to noise (see Fig. 3C). False alarms in this model can be prevented by setting a higher evidence criterion, i.e. creating a stricter reference point distinguishing target evidence samples from noise.

While both neurally-informed models mimic the behavioural patterns well (See Fig. 3B and C and Goodness of Fit in Table 1), the leak-adjustment model provides a poorer fit to the behavioural data than the criterion-adjustment model (G^2^ = 23; see Fig. 3B lower). This is largely due to the fact that the leak-adjustment model overestimates the number of False Alarms in the ITI, specifically in the Strong context. Remarkably, despite being constrained to adopt the precise bound settings indexed by Beta excursion, which oppose the adjustments predicted by the bound-adjustment model, the criterion-adjustment model was able to attain the same goodness-of-fit to behaviour as the initial bound-adjustment model (G^2^ = 16). To provide further model validation, the empirical CPP was compared against the simulated evidence-accumulation time courses of the different models. In the case of the neurally-informed models, predicted signatures of bound settings (e.g. cumulative evidence at response) are not informative for validation because they are forced to equal the observed Beta excursions by design (see discussion for comparison of the observed CPP pre-response amplitudes with observed Beta excursions). Instead, for the comparison of these two models, respectively featuring leaky accumulation and criterion-referenced, non-leaky accumulation, the stimulus-locked dynamics of accumulator buildup rate (i.e. first derivative) are particularly informative. While a leaky accumulator would be expected to begin to decline in its steepness immediately as it starts to build, a non-leaky accumulator would be expected to rise linearly with a steady slope for a certain amount of time. To test this and compare the models, we analysed the slope profile of the observed and simulated accumulation waveforms. We focused on the Strong conditions which will be the most diagnostic in this regard because the strong evidence onset produces a high, robust initial slope, as well as a sharp and relatively invariant CPP buildup onset which best approximates the onset invariance assumed in the models. The Fixed Strong and, to a lesser degree, the Mixed conditions also had stronger predicted degrees of leakage, so that any leak-induced drop in slope should be most apparent against EEG noise. By comparison, the Weak conditions have a much less clear onset time presumably due to greater variability in buildup onset timing not accounted for in the model. We found that the empirical CPP slope time course plateaued in the Strong conditions indicating a linear buildup profile (Fig. 3D), whereas the simulated time course of the leaky accumulator model indeed showed a steep drop-off in temporal slope after an initial peak (Fig. 3E). Meanwhile, the criterion-adjustment model with non-leaky accumulation showed a plateau in the slope similar to the empirical CPP (Fig. 3F). To capture this key feature quantitatively, we computed the percentage drop in slope in the first 100 ms relative to its initial peak. For the Fixed Strong condition the empirical slope dropped by 5% (bootstrap CI_95_% = [-5% 16%]), while the criterion-adjustment model predicted a drop of 6%, and the leak-adjustment model a drop of 27.5%, the latter falling outside the empirical bootstrapped 95% CI. Similarly, in the Mixed Strong condition the empirical slope dropped by 1.5% (bootstrap CI_95_% = [-9.5% 13%]), compared to 3% predicted by the criterion-adjustment model and 17% predicted by the leak-adjustment model.

### Baseline and pre-response Beta amplitudes and their relationship with RT

The above analysis of the Beta excursion showed that participants began their decision process with a smaller distance to the motor execution threshold in the Strong Context than for the other conditions (Fig. 2A). The dynamics of Beta amplitude relative to the action-triggering threshold were the key element for constraining the models as these amplitudes dictate the criterion amount of cumulative evidence required to trigger a response. However, to help interpret these changes in terms of context-related differences in psychological state, it is also of interest to separately examine variations in absolute pre-evidence and pre-response beta amplitudes across and within conditions. At first blush the smaller excursion in the Strong context appears consistent with participants being tonicially more prepared to make a movement, which seems counterintuitive for a task condition that is by far the easiest. However, pre-evidence Beta amplitude was actually found to be higher, reflecting lower baseline motor preparation in the Strong Context than both the Weak and Mixed Context (see Figures 4A left and B, *GMM, χ*^2^(1) = 2.3, p = 0.02 and *χ*^2^(1) =2.4, p = 0.02, respectively). Meanwhile, pre-response Beta amplitude was also significantly higher for the Strong Context, but to an even greater degree than at baseline (see Figures 4A right and C, *GMM, χ*^2^(1) = 3, p = 0.003). Thus, compared to the Mixed and Weak contexts, the much easier Strong context appeared to have lower tonic levels of preparation, but also a disproportionately more liberal threshold level, resulting in the smaller Beta excursion overall.

**Figure 4.**
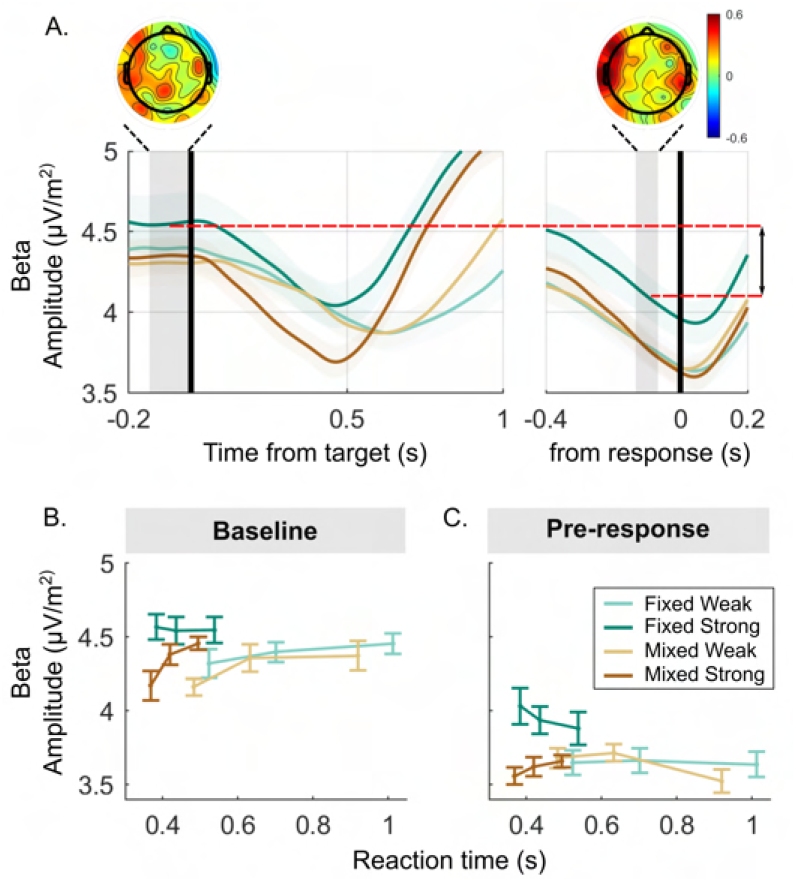
Detailed analysis of raw Beta amplitude in the baseline and pre-response timeframes. **(A)** Beta amplitude (15-30 Hz) over the motor cortex without any baseline- or threshold-subtraction, aligned to the target (*left)* and to the response (*right*), averaged for each condition. Additionally, a visual representation of excursion is shown. Scalp topographies show the distribution of the difference between strong targets in the Mixed and Fixed condition, red indicating higher activity in the Fixed condition. **(B)** Baseline Beta activity and **(C)** pre-response Beta plotted for each condition and three equal-size RT bins. Error bars for all plots represent the 95% confidence interval.

We also examined variation of pre-target Beta amplitude as a function of RT. As stated above, Beta excursion was larger on trials with longer RTs. Interestingly, examining the baseline and pre-response amplitudes separately indicated that this RT relationship was driven by a correlation with Beta amplitude in the baseline for the Mixed and Weak conditions (as seen previously, e.g. ***Steinemann et al., 2018***), whereas it was driven by a correlation with pre-response amplitude in the Strong context. This was indicated by a significant interaction between Context, Motion coherence, and RT for both baseline Beta (*GMM*, *χ*^2^(1) = −0.43, p = 0.004) and pre-response Beta (*GMM*, *χ*^2^(1) = −0.3, p = 0.007). Post-hoc tests showed that baseline Beta significantly varied with RT for both Weak conditions (Both Weak conditions p < 0.02) and the Mixed Strong conditions (*GMM*, *χ*^2^(1) = −0.32, p < 0.001), but not Fixed Strong (Fig. **4**B; *GMM, χ*^2^(1) = −0.009, p = 0.93). In contrast, pre-response Beta predicts RT in only the Fixed Strong condition (Fig. **4**C; *GMM, χ*^2^(1) = 2.5, p = 0.02 vs. Weak: p > 0.75 and Mixed Strong: *χ*^2^(1) = −0.05, p = 0.52). Thus, while excursion predicted RT in all conditions, this relationship with RT was expressed differently in the absolute baseline and pre-response levels in the Fixed Strong context, potential reasons for which are discussed below.

## Discussion

In order to optimise detection task performance, which requires balancing true (signal-driven) against false (noise-driven) detections, the decision process must be strategically adjusted according to various task factors including one’s knowledge of the evidence strength compared to the noise. In classic signal detection theory (SDT), this is accomplished by adjusting a decision criterion which the observer uses to categorise a unitary observation as either signal or noise, setting it lower (more liberal) when targets are known to be weak. However, the normative SDT framework is based on a unitary sensory observation. In continuous detection scenarios, where fast responses are required to unpredictable targets within ongoing noisy information, decisions benefit from accumulating evidence over time. An obvious analogous mechanism for setting liberal/conservative policies in this case is to adjust a criterion on cumulative evidence, in the form of a response-triggering decision bound, as has been found to regulate speed-accuracy tradeoff in discrete forced-choice decisions (***Churchland et al., 2008***; ***Hanks et al., 2014***; ***Murphy et al., 2016***; ***Palmer et al., 2005***; ***Reddi and Carpenter, 2000***; ***Reddi et al., 2003***; ***Thura and Cisek, 2016***). Here we tested this prediction through the joint analysis and modelling of behavioural and electrophysiological data in a continuous motion detection task performed in three different difficulty contexts. Although we confirmed that contextdependent behaviour can be explained by a bound-adjustment model, we found that Beta activity over motor cortex, a well-established neural signature of a thresholded decision variable, showed adjustment trends in the opposite direction. Beta excursion from baseline to motor execution threshold was smaller in the Fixed Strong context, indicative of a more liberal bound in this condition (see Figures 2A and B). Using the exact urgency dynamics reflected in Beta oscillations which dictate the decision variable’s starting level relative to threshold and hence the decision bound, we showed that behaviour can be captured equally well by a continuous accumulation model that employs an adjustable sensory criterion to which momentary evidence is referenced before accumulation.

Building on other recent studies using neurally-informed modelling (reviewed in ***O’Connell and Kelly, 2021***), the current study highlights the value in incorporating neural signatures of decision formation in models to uncover effects that cannot be resolved by considering behaviour alone. Specifically, the discrepancy between the behaviour and Beta excursion proved to be crucial to test alternative computational mechanisms to implement context-adjustments. The essential characteristic of the sensory-criterion adjustment mechanism is that it controls the strength of sensory input that is sufficient to give rise to significant positive accumulation. Within the framework of the current study, in which we assumed a common basis of continuous accumulation for all models, we implemented the criterion as a reference-point subtracted from all evidence samples as they enter a continuous accumulator with lower reflecting bound. However, future research should systematically explore other viable implementations of the sensory criterion mechanism. In particular, accumulation need not be continuous and could instead be triggered, for example by the evidence first reaching a “gate” level as has been proposed for visual search decisions (***Purcell et al., 2010, 2012***; ***Schall et al., 2011***), or by a distinct, lower-level process functioning as a relevant transition detector, neural correlates of which have been reported in humans (***Loughnane et al., 2016***) and are also present in the current data (see Fig. 2 - figure supplement 1). These gate-based triggering mechanisms provide alternative ways to implement an adjustable criterion to regulate the input to the accumulator in order to set decision policy, and could potentially be teased apart through more complex manipulations of noise and target evidence dynamics.

In previous studies it was found that Beta activity reached a stereotyped pre-response amplitude contralateral to the response across different conditions and RTs, suggesting a fixed threshold applied to motor preparation (***O’Connell et al., 2012***; ***Steinemann et al., 2018***; ***Kelly et al., 2021***). While this holds true for most conditions in the current data, pre-response Beta activity in the Fixed Strong condition was significantly elevated relative to the other conditions (see Fig. 4A). We side-stepped this issue in our models by constraining them using the excursion relative to the pre-response level in a given condition rather than the absolute baseline activity. However, follow-up analysis of absolute Beta levels yields further insight into how the context-dependent adjustments are physiologically implemented. Although the smaller Beta excursion in the Fixed Strong condition might suggest that people maintained a higher state of continuous preparedness, which is presumably energy-consuming (***Gottsdanker, 1975***; ***Näätänen, 1972***) and thus counterintuitive for such an easy task. In fact, the elevated baseline Beta activity suggests that participants could afford to adopt minimal tonic preparation in this context. Meanwhile the yet greater elevation in pre-response Beta could be interpreted as a more relaxed level of response caution at the motor level, which in our criterion-adjustment model is counteracted by the higher sensory-level criterion to create an overall more conservative policy. We also found that pre-response Beta amplitude in the Fixed Strong condition was higher for faster RTs than for slower RTs. However, relative to the Mixed Strong condition, this elevation for the fastest trials was also observed in the baseline period (Fig. 4B), suggesting that other, slower- fluctuating generators of Beta activity that also relate to RT may superimpose on the motor preparation signal. Indeed, Beta activity has been linked to many cognitive factors other than motor preparation (***Aoki et al., 2001***; ***Brovelli et al., 2004***; ***Buschman and Miller, 2007, 2010***; ***Engel and Fries, 2010***; ***Murthy and Fetz, 1992***; ***Siegel et al., 2012***; ***Spitzer, 2017***), and here the topographies of condition differences suggest context-dependent changes across a wider network (Fig. 4A). This again underlines the importance of using beta excursion as model constraints to minimise the influence of slower-fluctuating interfering Beta generators, but also highlights the need for future research into how other factors such as attention, response vigour and sensorimotor connectivity can influence pre-response contralateral Beta amplitudes.

An advantage of having characterised multiple neural signatures of decision formation is that one can be used to constrain models while the other can be used for additional follow-up validation. However, in these data the observed CPP and Beta signals were not precisely aligned in their respective estimates of decision bound settings. Specifically, the pre-response amplitude of the CPP (Fig. 2E) failed to mirror the reduction in Beta excursion seen in the Fixed Strong condition (Fig. 2B). Under the assumption that the CPP reflects the pure cumulative evidence component that adds to urgency at the downstream motor level where the threshold is set, it should reach lower pre-response amplitudes for conditions in which Beta excursion is smaller (***Steinemann et al., 2018***; ***Kelly et al., 2021***). However, simulations of both neurally-informed models (see Fig. 3 - figure supplement 1C and D) confirm that the predicted reduction in pre-response accumulator amplitude for the Fixed Strong relative to Mixed Strong is very small (6%), and not an effect easily detected in event-related potentials, suggesting the possibility of a type 2 error. Nevertheless, it is possible that the true empirical CPP amplitude in the Fixed Strong condition exceeds the value predicted in the models. we could speculate that this could arise from a relaxation of either the weighting or the delay of the transmission of CPP information into motor preparation in this easy condition, either of which would cause the amplitudes to be higher in the fixed pre-response window without changes to model structure or constraints.

The sensory-criterion adjustment mechanism presents an interesting alternative means of policy-setting with respect to the strategic adjustment of the action-triggering bound suggested by the dominant computational models and empirical research on the speed-accuracy tradeoff (***Bogacz et al., 2010***; ***Palmer et al., 2005***). It is important to note, however, that it is not yet clear why the policy is set by sensory criterion setting here and by decision bound setting in the speed/accuracy work, as there are two major differences: here the decision requires an adjustment for sensory target strength not speed/accuracy emphasis, and here the task is detection within a continuous stimulus as opposed to discrimination within discrete trials. To tell which factor explains the difference in adjustment mechanism, it will be necessary to examine speed-accuracy tradeoff manipulations in a continuous detection task.

In conclusion, while previous studies of decision policy adjustments for other perceptual task manipulations have found that they are enacted by changing the decision bound, we here show that, when adjusting for expected difficulty level during continuous detection, people rather regulate the transfer of sensory evidence into the accumulator process. This shows that the brain harbours a broader range of mechanisms to regulate decision policy.

## Methods and Materials

### Participants

Fourteen participants (aged 21-60 years; 8 female) were recruited at University College Dublin. Two participants were left-handed, all participants had normal or corrected-to-normal visual acuity, and had no previously diagnosed psychiatric disorders, epilepsy or suffered from any head injury resulting in loss of consciousness. All participants gave written consent prior to their participation and were compensated for their time with €25. The UCD Human Research Ethics Committee for Life Sciences approved all experimental procedures in accordance with the Declaration of Helsinki.

### Experimental procedure

Participants performed the task in a dark room, while seated 57 cm from the monitor. All stimuli were presented on a black background on a 21-inch CRT monitor operating at 60 Hz refresh rate (1280 x 960 pixels). Stimuli were generated using custom-made software in Matlab®R2016 (Mathworks®, Inc., Natick, MA, USA) utilising Psychtoolbox (***Brainard, 1997***; ***Kleiner et al., 2007***; ***Pelli, 1997***).

#### Experimental tesk: Random Dot Motion

Participants performed a continuous detection version of the random dot motion task (RDM; ***Hanks et al., 2006***; ***Roitman and Shadlen, 2002***; ***Ditterich, 2006***; ***Shadlen and Newsome, 1996***). In this task, they were asked to continuously monitor a cloud of white, randomly moving dots for intermittent targets defined as a step change from random to coherent upwards dot motion lasting for 1 sec (see Fig. 1A). During periods of incoherent motion (0% motion coherence), all dots were randomly displaced to a new location throughout the patch on each frame. Coherent dot motion was accomplished by displaying a certain percentage of randomly selected dots in a direction relative to their previous location within each frame. The inter-target interval (ITI), in which motion coherence was held at 0%, was pseudo-randomly varied among 4 possibilities: 2, 4, 6, or 8 sec. This ITI variability created temporal uncertainty to ensure continuous engagement in the detection task. Three target-strength contexts were run in separate blocks: in the ‘Weak Evidence Context’ all targets had 25% coherence; in the ‘Strong Evidence Context’ all targets were 70%; and both coherence levels could appear with equal likelihood in a ‘Mixed Context’. Participants were asked to respond to coherent upwards motion by clicking a mouse button with the index finger of their dominant hand. Each participant performed 12 blocks each consisting of 24 targets. Three blocks of each of the “Fixed” (single-coherence targets) contexts (“Strong” or “Weak”) were first run in an order counterbalanced across participants. The “Mixed” target coherence context was originally designed as a control condition to verify bound invariance across targets’ strengths within a single context, and thus all participants performed six “Mixed” blocks at the end of the experimental session.

Prior to the experiment, participants completed four training blocks, each consisting of 6 targets with decreasing levels of coherence (90%, 70%, 45%, 25%). In the last training block, participants would practise the Mixed Context condition (25% and 70%). Feedback of the participant’s performance was presented in text at the end of each block (numbers of hits, misses, and false alarms). The RDM pattern consists of a patch of 150 white, randomly moving dots (dot size: 4 pixels, dot speed of 3.33 deg/s) centrally presented in an aperture of 8° diameter. Dots were flickering on and off at a rate of 15 Hz. While this flicker was included as standard at the time to measure steady-state visual evoked potentials (see also ***Kelly and O’Connell, 2013***), these potentials do not reflect sensory evidence for the decision (which is instead the motion coherence) and therefore were not analysed here.

### Data analysis

#### Behavioural analysis

All trials were sorted and analysed according to the four conditions of Fixed Weak, Fixed Strong, Mixed Weak and Mixed Strong. Reaction time (RT) was calculated in milliseconds (ms) relative to the evidence (i.e. coherent dot motion) onset. Hits were defined as trials in which a response was made between 200 ms after target-onset up to 650 ms after target-offset, by the logic that responses with RT faster than 200 ms are defined as false alarms (e.g. uninformed by the sensory evidence) and responses 650 ms after target offset are defined as late responses (this was the point where the response probability returned to false alarm rate). Conversely, false alarms were defined as responses in the period from 650 ms after target-offset to 200 ms after the following target onset. False alarm rate was calculated as the proportion of 2-sec periods of inter-target interval time in which such a false alarm response was made. We chose this “per 2-sec” scaling as it is the shortest ITI but note that no results are dependent on this choice of scaling, as it merely sets the units of measurement.

Effects of the different conditions on RT were statistically analysed using a repeated-measures ANOVA (rANOVA) that organised the 4 conditions into two binary factors of Difficulty context (Fixed vs. Mixed), and Coherence (25% vs. 70%), and with the additional factor of ITI duration preceding a given target (2, 4, 6, or 8 s). Generalised mixed regression models (GMM) were used to analyse hit rate and false alarms rate, which allows statistical testing of proportions and bounded variables. A logit link-function was used for hit rates as they are binomial distributed. False alarm rates were analysed as a function of Context (Weak, Strong, and Mixed Contexts). A log-link function is used for the false alarm rates as participants had unbounded opportunities to make false alarms within each ITI, yielding a poisson distribution. For all models, intercepts and slopes were added as random-effects.

#### EEG acquisition and preprocessing

Continuous EEG was recorded from 128 scalp electrodes using a BioSemi system with a sample rate of 512 Hz. All data was analysed in a Matlab R2018a utilising EEGLAB routines (Delorme, et al., 2004). Eye movements were recorded with four electro-oculogram (EOG) electrodes, two above and below the left eye and two at the outer canthus of each for vertical (vEOG) and horizontal (hEOG) eye movements and blinks. EEG data were low-pass filtered by convolution with a 137-tap hanning windowed sinc function to give a 3-dB corner frequency of 37 Hz with strong attenuation at the mains frequency (50 Hz; ***Kelly et al., 2021***; ***Widmann et al., 2015***). Noisy channels were automatically detected using the PREP pipeline (***Bigdely-Shamlo et al., 2015***) and interpolated using spherical spline interpolation. Initially, large target epochs were extracted using a window capturing the preceding ITI (up to 650 ms after the previous target-offset) to 1 sec after the current target-offset.

Trails were rejected if the absolute difference between the vEOG or hEOG electrodes exceeded 200 *μ*V during −100 ms before target-onset up to 100 ms after response or if 10% of the electrodes exceeded 100 *μ*V in the target epoch. If less than 10% of the electrodes exceeded 80 *μ*V on a given trial, then these electrodes were interpolated using a spherical spline interpolation. Additionally, trials with a RT slower than 1650 ms were rejected. One participant was excluded due to excessive trial rejection (>70% in each condition) due to a mix of blinking, eye movements and EEG artefacts. After artefact rejection, the data were subjected to a current source density (CSD) transformation utilising the CSD toolbox in Matlab®R2018a (***Kayser and Tenke, 2006***; ***Kayser, 2009***).

#### Decision signal analysis

We analysed two neurophysiological signals previously established to reflect decision formation. First, 1) effector-selective spectral amplitude decreases in the Beta (15-30 Hz) frequency-range amplitude reflecting motor preparation. We omitted the 15 Hz frequency bin to avoid contamination of more preparation signals by the 15-Hz steady state visually evoked potentials (SSVEP) evoked by the on-off flicker of the dots at the same frequency. Second, we analysed 2) the CPP, a motor-independent signal exhibiting evidence accumulation dynamics (***O’Connell et al., 2012***; ***Kelly and O’Connell, 2013***). To measure Beta amplitude, we computed a Short Time Fourier Transform (STFT) by applying a FFT in a sliding window of 266 ms (4 full cycles of the SSVEP) in steps of 10 ms, and then extracted amplitude in the 15 - 30 Hz frequency range for the electrode at standard sensorimotor sites C3 and C4, for participants that responded with right and left hand, respectively.

The CPP was measured from three centro-parietal electrodes that we selected from each individual participant with the highest signal to noise ratio (amplitude −133 to −66 ms before the response divided by the standard measurement error as suggested by ***Luck et al.*** (***2021***)), within a cluster of nine electrodes delineated based on the visual inspection of the grand average topography of the response CPP (see topography in Fig. 2D). Beta and CPP signals were then segmented into target-locked and response-locked epochs. Target-locked epochs were extracted from −200 ms to 1000 ms around target-onset, and response-locked epochs were extracted from −400 to 200 ms around response time. For the CPP, epochs were baseline-corrected to −133 to −66 ms around target-onset (two full cycles of the SSVEP at 15 Hz). They were additionally low-passed filtered at 6 Hz with a fourth-order Butterworth filter, for plotting purposes only.

To determine whether the neural signals show signs of decision bound adjustment, we compared the Beta amplitude “excursion” — i.e., amplitude just prior to response relative to just prior to target onset— and the pre-response CPP amplitude across the four conditions. Beta Excursion was calculated on each trial by taking the difference of mean baseline amplitude (−133 to −66 ms relative to target-onset) minus mean pre-response amplitude (−133 to −66 ms relative to response). CPP amplitude was calculated for each trial as the average amplitude in the pre-response time range −133 to −66 ms relative to response. The effects on Beta Excursion and CPP were examined using a GMM with the factors of Difficulty context (Fixed vs. Mixed) and Motion coherence (25% vs. 70%), and using RT (z-scored within participant) as a covariant factor. For all models, random intercepts and slopes were added as random-effects.

As there was an increase of false alarms as the ITI progressed, we examined evidence-independent urgency by analysing grand-average Beta amplitude in the 8-sec ITI for the Strong, Weak and Mixed contexts. In order to create a standardised urgency signal to constrain computational decision models (see below), we took the grand-average beta amplitude for each context in the pre-response amplitude (−67 to −33 ms) as the threshold level for that context and subtracted it from the beta signals measured during the ITI. Subsequently, we fitted a 3rd-order polynomial to capture the smooth urgency trend underlying the signal, ignoring any residual noise. The smooth urgency function was then normalised by dividing it by the maximum value during the ITI in any context, and finally it was inverted (urgency = 1-urgency). In this way, zero corresponds to the lowest motor preparation level measured during the ITI in the grand average Beta signals of any context and 1 corresponds to the threshold level for triggering a response.

#### Computational models

We fitted three alternative models to the behavioural data: a leaky accumulator model with context-dependent bound (henceforth the Bound-adjustment model) and two neurally-constrained models with urgency quantified directly from Beta signals as above, and featuring either leaky accumulation with context-dependent leak (henceforth the Leak-adjustment model), or non-leaky accumulation of evidence cast relative to a sensory criterion with a lower reflecting decision bound at zero (henceforth the Criterion-adjustment model). All three models are based following basic leaky-accumulator model equation (***Ossmy et al., 2013***):

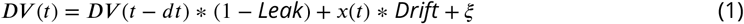

where *x*(*t*) is a binary representation of target-presence throughout the block,(set to 1 during targets and 0 otherwise), which is scaled by drift rate “Drift” according to the strength of the evidence for a given target and with added Gaussian noise, 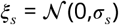, altogether representing the noisy sensory evidence through the block. As indicated in the equation, the *Leak* parameter sets the degree to which the previous accumulator value is scaled-down relative to the incoming evidence. When the accumulator level (*DV(t)* exceeded the decision-bound (*θ*), it was then set to zero for a post-response pause of 1 second, on the assumption that subjects are unlikely to resume accumulation immediately after responding given the range of ITIs.

Additionally, a non-decision time accounting for the delay between samples of evidence appearing on the screen and registering as increments to the accumulation process was estimated as a free parameter, while post-commitment delays associated with motor execution were fixed at 80 ms based on a typical motor time estimated for mouse-button presses (***O’Connell et al., 2012***; ***Steinemann et al., 2018***; ***Kelly et al., 2021***). The apportioning of non-decision delays to before or after the accumulation to bound process does not generally impact the fitting of behavioural models, but we applied this fixed motor time here to be consistent with the measurement time-window used for CPP amplitude. For the purposes of comparing the simulated and observed CPPs, rather than setting DV=0 immediately upon threshold crossing we allowed the simulated accumulator to continue accumulating for 80 ms beyond the point of decision commitment, after which the accumulator linearly decreased back to 0 over a period of 400 ms, mimicking the overshoot and fall of the empirical CPP after the response time. Note that model fits to behaviour are unaffected by the inclusion of this post-decision ramp down. Further, to ensure that comparison of real and simulated CPP time courses would be minimally influenced by any misestimation of such time-shifts, we took measurements relative to the signal onset time in that analysis.

#### Bound-adjustment model

A leaky integrator model (Fig. 1C), where the decision bound was free to vary across the three contexts (Weak, Strong and Mixed), was fitted only to the behavioural data as the model predicted a priori to explain the decision process adaptations. This model fixed the momentary sensory noise parameter at a value of 0.1 (*σ_s_* = 0.1), and included three free decision-bound parameters —one for each context (*θ_context_*)—, one *Leak* parameter, commonly applied to all conditions, one non-decision time parameter, and one or four *Drift* rate parameters (see below), making a total of 6 or 9 free parameters.

#### Neurally-constrained leak-adjustment model

Using the same leaky accumulator mechanism as above, this model (Fig. 3B) instead three leak parameters (*Leak_context_*) so that leak was free to vary across the contexts (Weak, Strong and Mixed). Here, the decision bound was fixed to 1 and additionally the sensory noise parameter (*σ_s_*) was a free parameter since the decision variable is now scaled by the normalised bound. Here, the sum *DV(t) + u(t)* was subjected to the fixed decision bound to determine responses (with the non-decision time implemented as the in the previous model), where u(t) is the urgency function measured from observed beta signals as detailed above in Decision Signal Analysis. This resulted in six free parameters for the minimal model with one free *Drift* rate parameter, and 9 for the expanded model with 4 free *Drift* rates.

#### Neurally-constrained criterion-adjustment model

In a second neurally-constrained model (Fig. 3C), we considered a non-leaky accumulation of evidence that is recast as a quantity relative to a sensory-level criterion (as distinct from a criterion set on cumulative evidence in the decision variable, which we refer to as a ‘decision bound’), with the additional; setting of a lower reflecting bound at zero on the accumulator, according to:

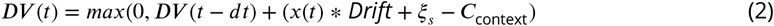

Here, sensory criterion (*C_context_*) was free to vary across the three contexts (Weak, Strong and Mixed). The fixed bound, constrained urgency and free noise parameter are exactly as in the neurally-constrained leak-adjustment model above. Again, the most minimal model with a single free drift rate parameter thus has a total of 6 free parameters (including one non-decision time as usual), and 9 free parameters for the version with 4 free *Drift* rates.

#### Behavioural model fitting

These models were each fitted to a total of 32 behavioural data points, comprised of the proportion of all targets with a hit in 6 RT bins defined by the .1, .3, .5, .7, and .9 RT quantiles or with no response (misses) across the four target conditions, and the false alarm rate measured as a proportion of every 2-second segment of the ITIs before the targets of each condition (see above and Fig. 1B). The models were fitted using Monte Carlo simulations of the decision process to generate simulated behavioural responses, which were compared to the observed data using the *χ*^2^ statistic *G*^2^:

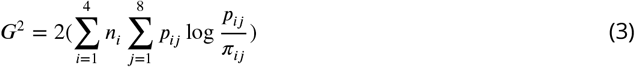

Here, *n_i_* is the number of valid trials, *p_ij_* and *π_ij_* are the observed and the simulated proportion, respectively, for condition *i* in RT bin *j. G*^2^ statistic was minimised with a bounded Nelder-Mead SIMPLEX algorithm (Nelder and Mead, 1965; fminsearchbnd in Matlab®).

Similar to ***Corbett et al.*** (***2021***), the models were fitted in several stages: In the first stage, a minimal model was fitted that included all relevant free parameters for that model (see above) but only one free drift rate parameter for the Weak Coherence, which was then linearly scaled for the Strong conditions according to relative coherence (70/25). In this initial broad search, about 4,500 trials were simulated per model evaluation in separate SIMPLEX fits for 1,000 different initial “guess” parameter vectors. These initial guess vectors were made by sampling from a uniform distribution spanning a reasonable range along each parameter dimension, so that the guess vectors together comprehensively covered a large parameter space in which to find a global minimum. In a second refinement stage, about 20,000 trials were simulated in each model evaluation in separate SIMPLEX fits starting from the 30 best estimated parameter vectors arising from the first step, with lower tolerance criteria for termination or convergence. This second step tended to improve the *G*^2^ only slightly, typically by about 2%.

Last, we ran a similar refinement step for model versions that now had four separate free drift rate parameters, to allow capturing any modulation of evidence accumulation quality due to either practice (as the participants always performed the mixed condition last) or context itself (e.g. drift rate may depend on how often strong targets arise in a given context). Using the 30 best refined parameters, we produced 200 new initial parameter vectors by adding random Gaussian jitter of a standard deviation of 1.1 times the mean of each of the refined parameters and 1.15 times of the mean for the drift rate parameters. Again about 20,000 trials were simulated in each model evaluation for this last step.

To confirm that the added complexity of four free drift rates was warranted over a single free drift rate parameter within each model, we computed the Akaike’s (AIC) and Bayes information on Criterion (BIC), to penalise for the complexity of the models, e.g. number of free parameters (*f*).

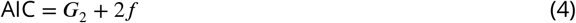

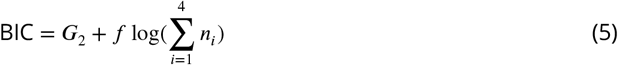

The principal comparisons in the study were across the three models with the same number of free parameters in any comparison, only differing in accumulation and/or adjustment mechanisms. Therefore these models were simply compared using *G*^2^ itself.

## Acknowledgments

S.P.K. and A.C.G. were supported by research grants from Science Foundation Ireland under grant number 15/CDA/3591 and The Wellcome Trust (219572/Z/19/Z). R.G.O. was supported by Horizon 2020 European Research Council Consolidator Grant IndDecision 865474.

---

**Figure 2—figure supplement 1.**
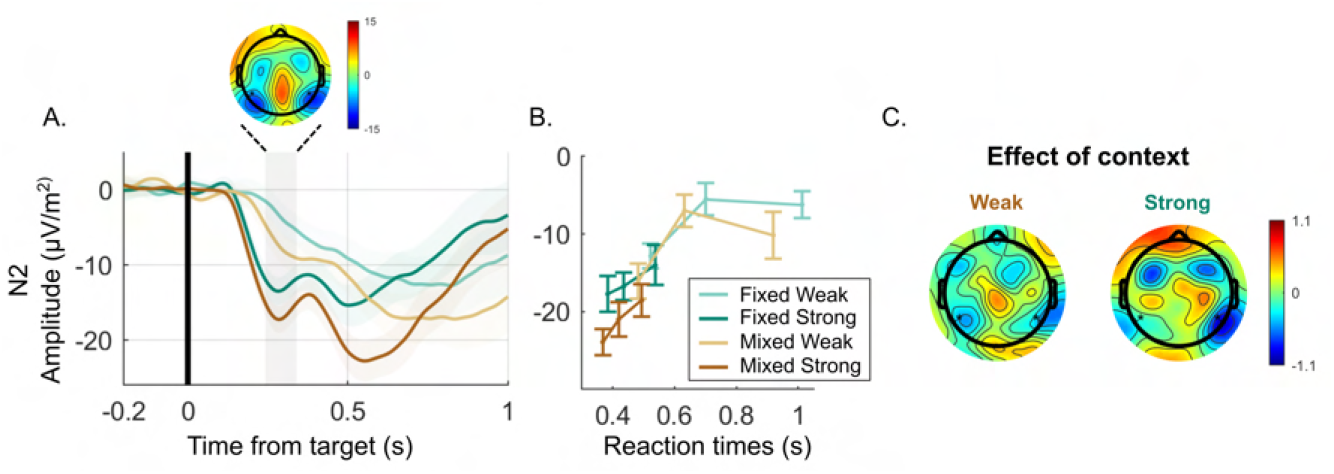
Posterior N2 component. **(A)** target-locked N2 **(B)** N2 amplitude around 300 ms after target onset, plotted as a function of RT for each condition. There are main effects of both target motion coherence (*χ*^2^(1) = 4.7, p = 0.006), with lower N2 for Strong targets as well as in the Mixed condition. Furthermore, there is a main effect of RT (*χ*^2^(1) = -2.6, p = 0.002), where faster RTs have lower N2s. **(C)** The scalp topography shows the distribution of the difference between Mixed and Fixed contexts. Blue would correspond with more negative amplitude in the Mixed relative to Fixed context.

**Figure 3—figure supplement 1.**
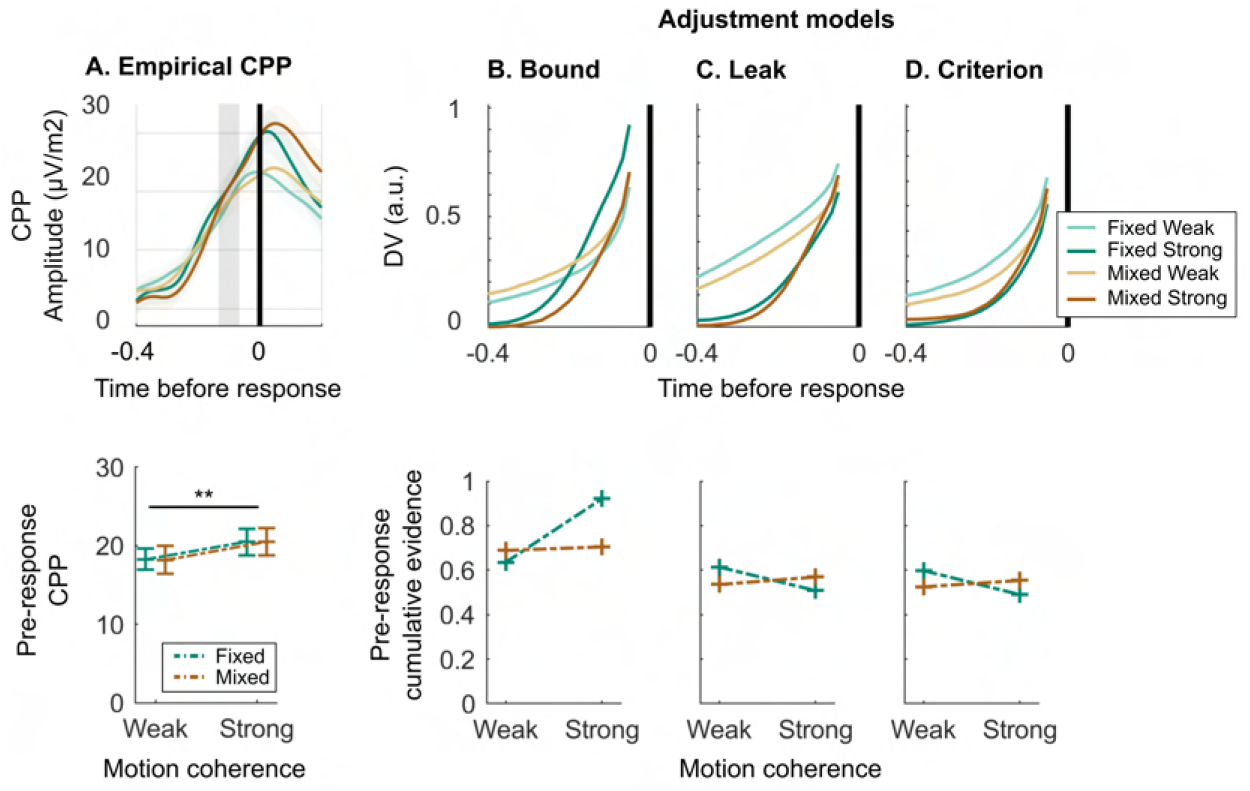
Comparison of response-locked dynamics in the empirical CPP. **(A)** to the simulated response-locked evidence accumulation dynamics of the bound-adjustment **(B)**, leak-adjustment **(C)** and criterion-adjustment **(D)** models. All 3 models predict an interaction in pre-response cumulative evidence levels, but whereas the bound adjustment model predicts the Fixed Strong condition to stand out as the highest, the neurally-informed models, by design, predict cumulative evidence levels that mirror the decision bounds estimated from empirically observed Beta excursion, where the Fixed Strong condition is instead the lowest.

## References

Afacan-Seref K, Steinemann NA, Blangero A, Kelly SP. Dynamic Interplay of Value and Sensory Information in High-Speed Decision Making. Current Biology. 2018; 28:795–802. https://doi.org/10.1016/j.cub.2018.01.071, doi: 10.1016/j.cub.2018.01.071.

Aoki F, Fetz EE, Shupe L, Lettich E, Ojemann GA. Changes in power and coherence of brain activity in human sensorimotor cortex during performance of visuomotor tasks. BioSystems. 2001; 63(1-3):89–99. doi: 10.1016/S0303-2647(01)00149-6.

Bigdely-Shamlo N, Mullen T, Kothe C, Su KM, Robbins KA. The PREP pipeline: Standardized preprocessing for large-scale EEG analysis. Frontiers in Neuroinformatics. 2015; 9(JUNE):1–19. doi: 10.3389/fninf.2015.00016.

Bogacz R, Wagenmakers EJ, Forstmann BU, Nieuwenhuis S. The neural basis of the speed-accuracy tradeoff. Trends in Neurosciences. 2010; 33(1):10–16. doi: 10.1016/j.tins.2009.09.002.

Brainard DH. The Psychophysics Toolbox. Spatial Vision. 1997; 10(4):433–6. doi: 10.1163/156856897X00357.

Brovelli A, Ding M, Ledberg A, Chen Y, Nakamura R, Bressler SL. Beta oscillations in a large-scale sensorimotor cortical network: Directional influences revealed by Granger causality. Proceedings of the National Academy of Sciences of the United States of America. 2004; 101(26):9849–9854. doi: 10.1073/pnas.0308538101.

Buschman TJ, Miller EK. Top-down versus bottom-up control of attention in the prefrontal and posterior parietal cortices. Science. 2007; 315(5820):1860–1864. doi: 10.1126/science.1138071.

Buschman TJ, Miller EK. Shifting the spotlight of attention: Evidence for discrete computations in cognition. Frontiers in Human Neuroscience. 2010; 4(November):1–9. doi: 10.3389/fnhum.2010.00194.

Churchland AK, Kiani R, Shadlen MN. Decision-making with multiple alternatives. Nature Neuroscience. 2008; 11(6):693–702. doi: 10.1038/nn.2123.

Cisek P, Puskas GA, El-Murr S. Decisions in changing conditions: The urgency-gating model. Journal of Neuroscience. 2009; 29(37):11560–71. doi: 10.1523/JNEUROSCI.1844-09.2009.

Corbett EA, Alexandra Martinez-Rodriguez L, Judd C, O’connell RG, Kelly SP. Multiphasic value biases in fast-paced decisions. bioRxiv. 2021; p. 2021.03.08.434248.https://doi.org/10.1101/2021.03.08.434248.

Ditterich J. Evidence for time-variant decision making. European Journal of Neuroscience. 2006; 24(12):3628–3641. doi: 10.1111/j.1460-9568.2006.05221.x.

Donner TH, Siegel M, Fries P, Engel AK. Buildup of Choice-Predictive Activity in Human Motor Cortex during Perceptual Decision Making. Current Biology. 2009; 19(18):1581–1585. http://dx.doi.org/10.1016/j.cub.2009.07.066, doi: 10.1016/j.cub.2009.07.066.

Donoghue JP, Sanes JN, Hatsopoulos NG, Gaál G. Neural discharge and local field potential oscillations in primate motor cortex during voluntary movements. Journal of Neurophysiology. 1998; 79(1):159–173. doi: 10.1152/jn.1998.79.1.159.

Engel AK, Fries P. Beta-band oscillations-signalling the status quo? Current Opinion in Neurobiology. 2010; 20(2):156–165. doi: 10.1016/j.conb.2010.02.015.

Forstmann BU, Anwander A, Schäfer A, Neumann J, Brown S, Wagenmakers EJ, Bogacz R, Turner R. Cortico-striatal connections predict control over speed and accuracy in perceptual decision making. Proceedings of the National Academy of Sciences of the United States of America. 2010; 107(36):15916–15920. doi: 10.1073/pnas.1004932107.

Forstmann BU, Dutilh G, Brown S, Neumann J, Von Cramon DY, Ridderinkhof KR, Wagenmakers EJ. Striatum and pre-SMA facilitate decision-making under time pressure. Proceedings of the National Academy of Sciences of the United States of America. 2008; 105(45):17538–17542. doi: 10.1073/pnas.0805903105.

Glaze CM, Kable JW, Gold JI. Normative evidence accumulation in unpredictable environments. eLife. 2015; 4(e08825):1–27. doi: 10.7554/eLife.08825.

Gold JI, Shadlen MN. The Neural Basis of Decision Making. Annual Review of Neuroscience. 2007; 30:535–74. doi: 10.1146/annurev.neuro.29.051605.113038.

Gold JI, Stocker AA. Visual Decision-Making in an Uncertain and Dynamic World. Annual Review of Vision Science. 2017; 3:227–250. doi: 10.1146/annurev-vision-111815-114511.

Gottsdanker R. Attention and performance. In: Rabbitts P, Dornic S, editors. The attaining and maintaining of preparation. London: Academic press; 1975.p. 3–49.

Gould IC, Nobre AC, Wyart V, Rushworth MFS. Effects of decision variables and intraparietal stimulation on sensorimotor oscillatory activity in the human brain. Journal of Neuroscience. 2012; 32(40):13805–13818. doi: 10.1523/JNEUROSCI.2200-12.2012.

Hanks TD, Ditterich J, Shadlen MN. Microstimulation of macaque area LIP affects decision-making in a motion discrimination task. Nature Neuroscience. 2006; 9(5):682–9. doi: 10.1038/nn1683.

Hanks TD, Kiani R, Shadlen MN. A neural mechanism of speed-accuracy tradeoff in macaque area LIP. eLife. 2014; 3(e02260):1–17. doi: 10.7554/eLife.02260.

Heitz RP, Schall JD. Neural Mechanisms of Speed-Accuracy Tradeoff. Neuron. 2012; 76(3):616–628. http://dx.doi.org/10.1016/j.neuron.2012.08.030, doi: 10.1016/j.neuron.2012.08.030.

Ivanoff J, Branning P, Marois R. fMRI evidence for a dual process account of the speed-accuracy tradeoff in decision-making. PLoS ONE. 2008; 3(7). doi: 10.1371/journal.pone.0002635.

Kayser J, Current source density (CSD) interpolation using spherical splines - CSD Toolbox; 2009.

Kayser J, Tenke CE. Principal components analysis of Laplacian waveforms as a generic method for identifying ERP generator patterns: I. Evaluation with auditory oddball tasks. Clinical Neurophysiology. 2006; 117(2):348–368. doi: 10.1016/j.clinph.2005.08.034.

Kelly SP, Corbett EA, O’Connell RG. Neurocomputational mechanisms of prior-informed perceptual decision-making in humans. Nature Human Behaviour. 2021; 5(4):467–81. doi: 10.1038/s41562-020-00967-9.

Kelly SP, O’Connell RG. Internal and external influences on the rate of sensory evidence accumulation in the human brain. Journal of Neuroscience. 2013; 33(50):19434–41. doi: 10.1523/JNEUROSCI.3355-13.2013.

Kleiner M, Brainard D, Pelli D, Ingling A, Murray R, Broussard C. What’s new in psychtoolbox-3. Perception. 2007; 36(14):1–16.

de Lange FP, Rahnev DA, Donner TH, Lau H. Prestimulus oscillatory activity over motor cortex reflects perceptual expectations. Journal of Neuroscience. 2013; 33(4):1400–1410. doi: 10.1523/JNEUROSCI.1094-12.2013.

Lasley DJ, Cohn T. Detection of a Luminance Increment: Effect of Temporal Uncertainty. Journal of the Optical Society of America. 1981; 71(7):845–850. doi: 10.1364/JOSA.71.000845.

Loughnane GM, Newman DP, Bellgrove MA, Lalor EC, Kelly SP, O’Connell RG. Target selection signals influence perceptual decisions by modulating the onset and rate of evidence accumulation. Current Biology. 2016; 26:496–502. http://dx.doi.org/10.1016/j.cub.2015.12.049, doi: 10.1016/j.cub.2015.12.049.

Luck SJ, Stewart AX, Simmons AM, Rhemtulla M. Standardized measurement error: A universal metric of data quality for averaged event-related potentials. Psychophysiology. 2021; 58(6):1–15. doi: 10.1111/psyp.13793.

Murphy PR, Boonstra E, Nieuwenhuis S. Global gain modulation generates time-dependent urgency during perceptual choice in humans. Nature Communications. 2016; 7(May):1–14. http://dx.doi.org/10.1038/ncomms13526, doi: 10.1038/ncomms13526.

Murphy PR, Wilming N, Hernandez-Bocanegra DC, Prat-Ortega G, Donner TH. Adaptive circuit dynamics across human cortex during evidence accumulation in changing environments. Nature Neuroscience. 2021; 24(7):987–997. http://dx.doi.org/10.1038/s41593-021-00839-z, doi: 10.1038/s41593-021-00839-z.

Murthy VN, Fetz EE. Coherent 25-To 35-Hz oscillations in the sensorimotor cortex of awake behaving monkeys. Proceedings of the National Academy of Sciences of the United States of America. 1992; 89(12):5670–5674. doi: 10.1073/pnas.89.12.5670.

Näätänen R. Time uncertainty and occurence uncertainty of the stimulus in a simple reaction time task. Acta Psychologica. 1972; 36(6):492–503. doi: 10.1016/0001-6918(72)90029-7.

O’Connell RG, Dockree PM, Kelly SP. A supramodal accumulation-to-bound signal that determines perceptual decisions in humans. Nature Neuroscience. 2012; 15(12):1729–35. doi: 10.1038/nn.3248.

O’Connell RG, Kelly SP. Neurophysiology of Human Perceptual Decision-Making. Annual Review of Neuroscience. 2021;44(1):495–516. doi: 10.1146/annurev-neuro-092019-100200.

O’Connell RG, Shadlen MN, Wong-Lin KF, Kelly SP. Bridging Neural and Computational Viewpoints on Perceptual Decision-Making. Trends in Neurosciences. 2018; 41(11):838–52. http://dx.doi.org/10.1016/j.tins.2018.06.005, doi: 10.1016/j.tins.2018.06.005.

Ossmy O, Moran R, Pfeffer T, Tsetsos K, Usher M, Donner TH. The timescale of perceptual evidence integration can be adapted to the environment. Current Biology. 2013; 23(11):981–986. http://dx.doi.org/10.1016/j.cub.2013.04.039, doi: 10.1016/j.cub.2013.04.039.

Palmer J, Huk AC, Shadlen MN. The effect of stimulus strength on the speed and accuracy of a perceptual decision. Journal of Vision. 2005; 5(5):376–404. doi: 10.1167/5.5.1.

Pelli DG, The VideoToolbox software for visual psychophysics: Transforming numbers into movies; 1997. doi: 10.1163/156856897X00366.

Pfurtscheller G, Stancák A, Neuper C. Post-movement beta synchronization. A correlate of an idling motor area? Electroencephalography and Clinical Neurophysiology. 1996; 98(4):281–293. doi: 10.1016/0013-4694(95)00258-8.

Purcell BA, Heitz RP, Cohen JY, Schall JD, Logan GD, Palmeri TJ. Neurally Constrained Modeling of Perceptual Decision Making. Psychological Review. 2010; 117(4):1113–1143. doi: 10.1037/a0020311.

Purcell BA, Schall JD, Logan GD, Palmeri TJ. From salience to saccades: Multiple-alternative gated stochastic accumulator model of visual search. Journal of Neuroscience. 2012; 32(10):3433–3446. doi: 10.1523/JNEUROSCI.4622-11.2012.

Ratcliff R. Group reaction time distributions and an analysis of distribution statistics. Psychological Bulletin. 1979; 86(3):446–461. doi: 10.1037/0033-2909.86.3.446.

Ratcliff R. Theoretical Interpretations of the Speed and Accuracy of Positive and Negative Responses. Psycho-logical Review. 1985; 92(2):212–25. doi: 10.1037/0033-295X.94.2.277.

Ratcliff R, Smith PL. A Comparison of Sequential Sampling Models for Two-Choice Reaction Time Roger Ratcliff Northwestern University Philip L. Smith University of Melbourne. Psychological review. 2004; 111(2):1 – 101. http://doi.apa.org/getdoi.cfm?doi=10.1037/0033-295X.111.2.333%0Apapers3://publication/doi/10.1037/0033-295X.111.2.333.

Ratcliff R, Tuerlinckx F. Estimating parameters of the diffusion model: Approaches to dealing with contaminant reaction times and parameter variability. Psychonomic Bulletin and Review. 2002; 9(3):438–481. doi: 10.3758/BF03196302.

Ratcliff R, Van Zandt T, McKoon G. Connectionist and diffusion models of reaction time. Psychological Review. 1999; 106(2):261–300. doi: 10.1037/0033-295X.106.2.261.

Reddi BAJ, Asrress KN, Carpenter RHS. Accuracy, Information, and Response Time in a Saccadic Decision Task. Journal of Neurophysiology. 2003; 90(5):3538–3546. doi: 10.1152/jn.00689.2002.

Reddi BAJ, Carpenter RHS. The influence of urgency on decision time. Nature Neuroscience. 2000; 3(8):827–830. doi: 10.1038/77739.

Roitman JD, Shadlen MN. Response of neurons in the lateral intraparietal area during a combined visual discrimination reaction time task. Journal of Neuroscience. 2002; 22(21):9475–9489. doi: 10.1523/jneurosci.22-21-09475.2002.

Schall JD, Purcell BA, Heitz RP, Logan GD, Palmeri TJ. Neural mechanisms of saccade target selection: Gated accumulator model of the visual-motor cascade. European Journal of Neuroscience. 2011; 33(11):1991–2002. doi: 10.1111/j.1460-9568.2011.07715.x.

Shadlen MN, Newsome WT. Motion perception: Seeing and deciding. Proceedings of the National Academy of Sciences of the United States of America. 1996; 93(2):628–633. doi: 10.1073/pnas.93.2.628.

Siegel M, Donner TH, Engel AK. Spectral fingerprints of large-scale neuronal interactions. Nature Reviews Neuroscience. 2012; 13(2):121–134. doi: 10.1038/nrn3137.

Spitzer B. Beyond the Status Quo : A Role for Beta Oscillations in Endogenous Content (Re) Activation. eNeuro. 2017; 4(4):1–15. doi: 10.1523/ENEURO.0170-17.2017.

Standage D, You H, Wang DH, Dorris MC. Gain modulation by an urgency signal controls the speed-accuracy trade-off in a network model of a cortical decision circuit. Frontiers in Computational Neuroscience. 2011; 5(February):1–14. doi: 10.3389/fncom.2011.00007.

Steinemann NA, O’Connell RG, Kelly SP. Decisions are expedited through multiple neural adjustments spanning the sensorimotor hierarchy. Nature Communications. 2018; 9(3627):1–13. http://dx.doi.org/10.1038/s41467-018-06117-0, doi: 10.1038/s41467-018-06117-0.

Thura D, Beauregard-Racine J, Fradet CW, Cisek P. Decision making by urgency gating: Theory and experimental support. Journal of Neurophysiology. 2012; 108:2912–30. doi: 10.1152/jn.01071.2011.

Thura D, Cisek P. Deliberation and commitment in the premotor and primary motor cortex during dynamic decision making. Neuron. 2014; 81(6):1401–1416. http://dx.doi.org/10.1016/j.neuron.2014.01.031, doi: 10.1016/j.neuron.2014.01.031.

Thura D, Cisek P. Modulation of premotor and primary motor cortical activity during volitional adjustments of speed-accuracy trade-offs. Journal of Neuroscience. 2016; 36(3):938–956. doi: 10.1523/JNEUROSCI.2230-15.2016.

Twomey DM, Murphy PR, Kelly SP, O’Connell RG. The classic P300 encodes a build-to-threshold decision variable. European Journal of Neuroscience. 2015; 42:1636–43. doi: 10.1111/ejn.12936.

Usher M, McClelland JL. The time course of perceptual choice: The leaky, competing accumulator model. Psychological Review. 2001; 108(3):550–92. doi: 10.1037/0033-295X.108.3.550.

Van Veen V, Krug MK, Carter CS. The Neural and Computational Basis of Controlled Speed-Accuracy Tradeoff The Neural and Computational Basis of Controlled Speed - Accuracy Tradeoff during Task Performance. 2Journal of Cognitive Neuroscience. 2008; 20(11):1952–65.

Wenzlaff H, Bauer M, Maess B, Heekeren HR. Neural characterization of the speed - Accuracy tradeoff in a per-ceptual decision-making task. Journal of Neuroscience. 2011; 31(4):1254–1266. doi: 10.1523/JNEUROSCI.4000-10.2011.

Widmann A, Schröger E, Maess B. Digital filter design for electrophysiological data - a practical approach. Journal of Neuroscience Methods. 2015; 250:34–46. doi: 10.1016/j.jneumeth.2014.08.002.

