## Supplementary figures and images for "Balancing true and false detection of intermittent sensory targets by adjusting the inputs to the evidence accumulation process"

### Figure 2 - supplementary 1

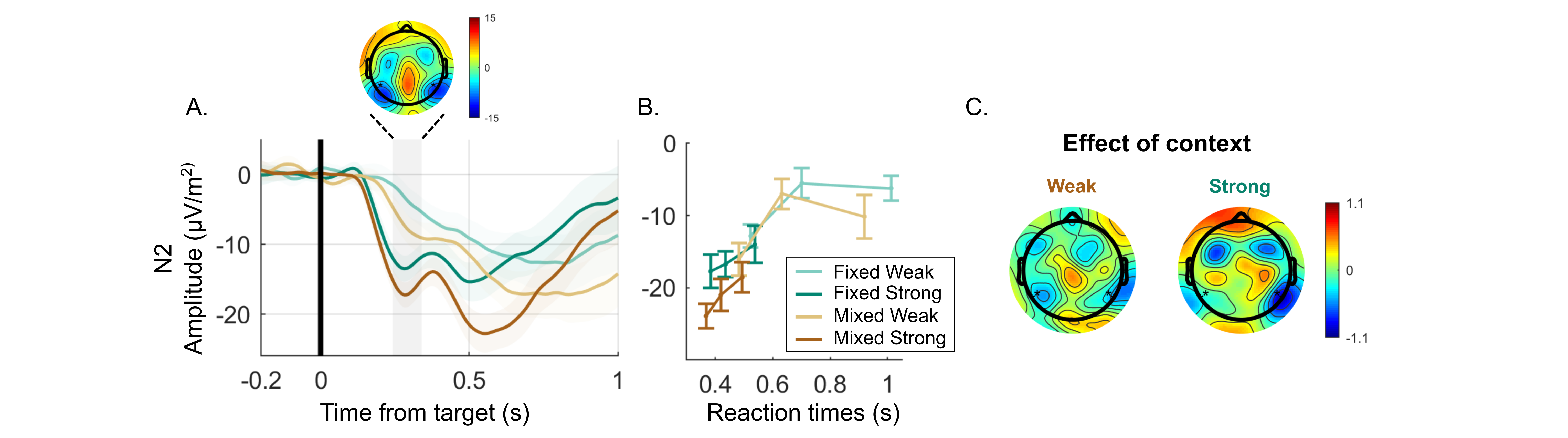

### Figure 3 - supplementary 1

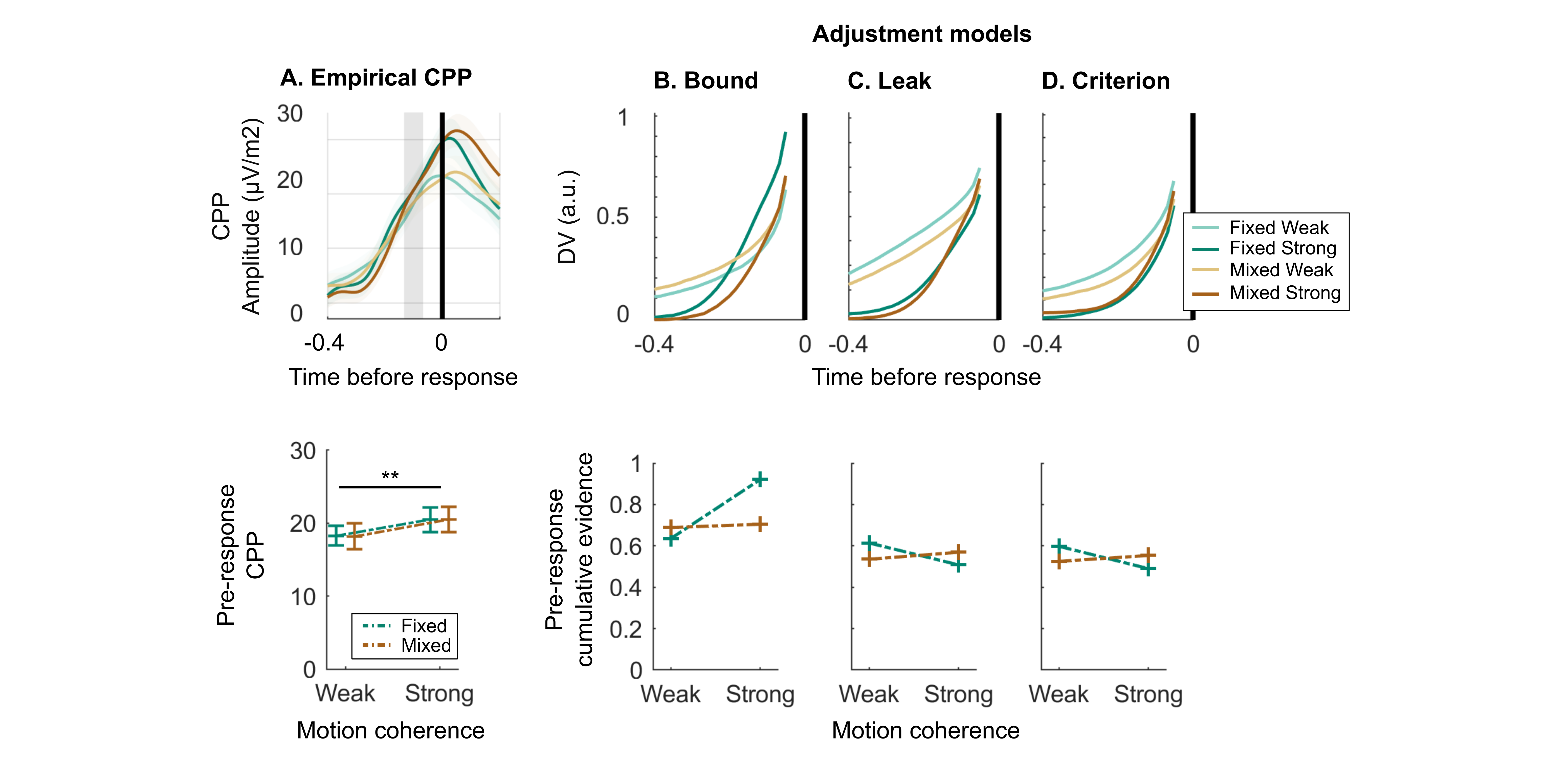
